# Validating a low-cost, open-source, locally manufactured workstation and computational pipeline for automated histopathology evaluation using deep learning

**DOI:** 10.1101/2023.04.19.537544

**Authors:** Divya Choudhury, James Dolezal, Emma Dyer, Sara Kochanny, Siddi Ramesh, Frederick M. Howard, Jayson R. Margalus, Amelia Schroeder, Jefree Schulte, Marina C. Garassino, Jakob N. Kather, Alexander T. Pearson

## Abstract

Deployment and access to state-of-the-art diagnostic technologies remains a fundamental challenge in providing equitable global cancer care to low-resource settings. The expansion of digital pathology in recent years and its interface with computational biomarkers provides an opportunity to democratize access to personalized medicine. Here we describe a low-cost platform for digital side capture and computational analysis composed of open-source components. The platform provides low-cost ($200) digital image capture from glass slides and is capable of real-time computational image analysis using an open-source deep learning (DL) algorithm and Raspberry Pi ($35) computer. We validate the performance of deep learning models’ performance using images captured from the open-source workstation and show similar model performance when compared against significantly more expensive standard institutional hardware.

## Introduction

The global burden of cancer is increasing as mortality associated with communicable diseases, starvation, and war declines. In the past a majority of cancer cases and cancer-related deaths occurred in higher income countries. However, the demographics of cancer are shifting: incidence of cancer in low Human Development Index countries is projected to double between 2008 and 2030 and increase by 81% in middle Human Development Index countries. Advances in cancer care disproportionately benefit people in high HDI countries: fewer cancer-related gains in life expectancy are seen in low versus high HDI countries [1, 2]. It is therefore crucial to adapt the complex cancer diagnostics currently used in high-resource settings for broader application. A hallmark of cancer diagnosis worldwide is pathological analysis with hematoxylin and eosin (H&E) stained tumor biopsy sections, and additional molecular tests can help aid treatment decision-making. Access to extensive molecular testing is currently limited by cost, so as the demographics of disease change and a growing number of people are diagnosed with cancers that require more complex diagnostic techniques, there is a growing need to close the diagnostic gap between high- and low-income regions.

Digital pathology involves the acquisition and analysis of digital histopathological images in place of conventional microscopy [3-5]. As digitized whole slide images (WSIs) of histopathologic slides become more widely available, computer vision and machine learning methods in digital pathology have the potential to automate diagnosis based on images of H&E slides. Deep learning (DL), which is a subdomain of machine learning that uses multi-layered convolutional neural networks to identify patterns and features in complex datasets [6, 7], can automate diagnostic workflows and reduce costs [8, 9] while still providing the same information that a human pathologist identifies in each histologic image [4, 10]. In addition, DL algorithms analyze higher-order image characteristics to identify important histologic and clinically actionable features including distinguishing cancerous from non-cancerous tissue, survival, treatment response, and genetic alterations [7, 11, 12].

Deep learning algorithms remain largely out of reach in low resource settings, where automation could offer an opportunity to address personnel shortages and lack of equipment as well as reduce costs associated with precision cancer care. Governments and corporations invest extensive resources into oncologic research to develop advanced diagnostics and targeted therapies, yet implementation barriers limit the ability of these advances to reduce the global burden of cancer. Specific advantages of digital pathology include the ability to share images in real time to allow for remote collaboration, easy acquisition of large amounts of data for analysis, and the potential to lower costs by reducing the amount of human pathologist review needed to make a diagnosis [5, 13]. These features make digital pathology well-suited for improving access to precision medicine in under-resourced areas.

However, state of the art DL-based methods for digital pathology diagnostics fail to integrate their technology with hardware and computing resources commonly found in lower-resource settings. As of November 2022, only one FDA-approved commercial AI technology for digital pathology exists, and no open-source technologies have gained approval [14, 15]. Purchasing commercial software, computing resources, and whole slide image scanners needed for currently available AI pathology tools costs hundreds of thousands of dollars, making them unusable in low resource settings. Additionally, validation of existing digital pathology DL methods has only been performed with high-cost equipment in high-income countries.

To overcome the cost barrier in digital pathology analysis, we compiled a fully integrated set of open-source, low-cost hardware and software components to compare DL model performance against high-cost methods. Several studies have shown that digital images from diagnostic glass slides can be captured with low-cost equipment at sufficient resolution for DL model analysis [16-20]. We constructed a workflow consisting of entirely open-source resources that integrates images captured with low-cost hardware, low-cost computing equipment, and publicly available DL models, thereby producing an end-to-end low-cost digital histology deep learning analysis toolkit [21]. We developed and tested a novel, $200 platform for image capture of histopathological slides and used an open-source DL pipeline run on a $35 Raspberry Pi computer to classify tissue samples from two distinct datasets. We show that model performance on images captured with low-cost equipment matches that of images captured with gold standard high-cost equipment and validated our pipeline for automated digital pathology biomarker-based classification of head and neck squamous cell carcinoma (HNSCC) and lung cancer subtypes.

## Materials and Methods

Data were collected to train DL models to perform various classification tasks. To predict human papillomavirus (HPV) status in HNSCC and lung cancer subtype, we collected publicly available histology images, which were captured using an Aperio ScanScope or other Aperio slide scanner and stored in SVS format from The Cancer Genome Atlas (TCGA). Tissue samples in these TCGA datasets originate from patients who live primarily in the United States and Europe [22]. Most patients’ primary tumor sites were classified as oral cavity tumors. To perform external validation on models trained with data from TCGA, we collected whole slide images (WSIs) from the University of Chicago Medical Center (UCMC), in accordance with University of Chicago IRB protocol 20-0238. For TCGA and UCMC datasets, pathologists identified and annotated tumor regions of interest (ROI) within each WSI. Patient demographics in the UCMC dataset were similar to those in the TCGA dataset.

### HPV status HNSCC dataset preparation

In this retrospective study, 472 digitized whole slide images (WSI) of H&E tissue samples from patients with HNSCC and a known HPV status were collected in SVS format from TCGA. Among this TCGA cohort, 52 patients were classified as HPV positive and 407 were classified as HPV negative. For HNSCC HPV status model testing, we used a UCMC external validation dataset consisting of ten histopathology glass slides collected from patients with HNSCC and a known HPV status. Of these patients, five had HPV positive and five had HPV negative cancers.

### Lung cancer dataset preparation

941 digitized whole slide images (WSI) of H&E tissue samples from patients with lung cancer of a known subtype were also collected in SVS format from TCGA. 472 tissue samples from this cohort were classified as lung adenocarcinoma and 469 were classified as squamous cell carcinoma. We also performed external validation of the lung classification model using a dataset from UCMC consisting of ten histopathology glass slides collected from patients with lung cancer of known subtype. Of these patients, five had lung adenocarcinoma and five had squamous cell carcinoma.

### Current standard imaging hardware and acquisition

For baseline “gold standard” validation image capture, we used a shared resource 3DHISTECH ($200,000) digital pathology microscope slide scanner for acquisition of images at 40X magnification. The 3DHISTECH scanner captures images in MRXS format.

### Open-source microscopy manufacture and image acquisition

To assess a cost-efficient alternative to professional grade microscopes, we used the OpenFlexure Microscope v6 design and assembled it with a fused deposition modeling 3D-printer (Creality CR-10s), which includes optics, illumination, and stage modules. Our current model of the device differs from the published OpenFlexure design in that it excludes the motorized stage. We used manual rather than motorized stage actuators to maximize stage mobility and attached the optics and illumination modules to the stage piece with the system of rails in the OpenFlexure design, securing the parts with m3 hex head screws. This microscope was then paired with a low-cost Raspberry Pi Model 4B ($35) and associated Camera Module 2 ($30) to acquire partial-slide images from ten UCMC HNSCC glass slides and ten UCMC lung cancer glass slides [21, 23].

Raw images captured at 10X magnification provided using the OpenFlexure device were generally clear, but most images had some degree of blur, color distortion, and/or spherical aberration. After manual calibration, effective optical magnification was determined to be approximately 0.4 microns per pixel. The Raspbian-OpenFlexure operating system provided the software needed to visualize the glass slide on a monitor as well as capture and save images. Images captured from the OpenFlexure device were saved in JPEG format on-device and used for subsequent analysis.

### Histology image processing

To assess the impact of low-vs high-cost image capture methods, we produced two sets of scanned histology images for the UCMC validation cohorts: 1) WSIs from a clinical-grade microscope, and 2) partial-slide images from the OpenFlexure device. Slides from the validation dataset were first scanned with the high-cost 3DHISTECH microscope, producing whole-slide images (WSIs) for each glass slide. For a subset of the slides in the validation studies, partial-slide images were then captured using the OpenFlexure microscope. As the OpenFlexure device is not yet capable of WSI capture, only partial slide images could be captured. For these slides, we captured predetermined Regions of Interest (ROIs) containing strongly predictive morphological features. This partial-slide, proof-of-concept data capture enabled us to demonstrate that histopathological images acquired using low-cost equipment can be accurately classified with open-source deep learning algorithms. We captured 100 images of the predetermined morphologically informative regions at 10X magnification using OpenFlexure software (OpenFlexure Connect) on the Raspberry Pi, five from each of the ten slides in the HPV validation dataset and five from each of the ten slides in the lung cancer validation dataset [21]. For both whole-slide (3DHISTECH) and partial-slide (OpenFlexure) images, DL predictions were generated on smaller image tiles (tile width 299 pixels and 302 μm), and final slide-level predictions were calculated by averaging tile predictions. All image processing was performed using Slideflow [24].

### Deep learning models

The HPV DL classification model was trained on the full HNSCC TCGA dataset to predict whether samples came from patients with HPV-positive or HPV-negative tumors. This model was then validated on the small institutional dataset from UCMC. The lung cancer classification model was trained to classify tumor tissue as either adenocarcinoma or squamous cell carcinoma on the full TCGA lung cancer dataset, and then validated on the UCMC institutional dataset. Weakly-supervised DL model training was performed using a convolutional neural network with an Xception-based architecture, ImageNet pretrained weights, and two hidden layers of width 1024 with dropout of 0.1 after each hidden layer [25]. Tiles received data augmentation with flipping, rotating, JPEG compression, and Gaussian blur. Model training was performed with 1 epoch and 3-fold cross-validation, using the Adam optimizer, a learning rate of 10^−4^. Models were trained using Slideflow version 1.2.3 [24] with the Tensorflow backend (version 2.8.2) in Python 3.8. All hyperparameters are listed in **Supplementary Table 1**.

### Computational resources

DL models require central processing units (CPUs) and graphical processing units (GPUs) with sufficient computational power to generate predictions. DL classification of the images captured using the 3DHISTECH microscope with Slideflow was carried out with an NVIDIA Titan RTX GPU ($3000) that is representative of the high-cost computational resources currently used for automated analysis of WSIs. To demonstrate the feasibility of using low-cost computational hardware to generate DL predictions, the images captured using the OpenFlexure device were classified using Slideflow hosted on a $35 Raspberry Pi 4B computer with a Broadcom BCM2711, Quad core Cortex-A72 (ARM v8) 64-bit SoC @ 1.5GHz processor and 4 GB of RAM.

### User interface

Slideflow includes an open-source graphical user interface (GUI) that allows for interactive visualization of model predictions from histopathological images (**Figure 2**). The Slideflow GUI can flexibly accommodate a variety of trained models for digital pathology image classification and can generate predictions from both Raspberry Pi camera capture, partial-slide images, and whole-slide images. We optimized this interface for deployment on low-memory, ARM-based edge devices including the Raspberry Pi and used the interface to generate predictions for all images captured on the OpenFlexure device.

**Figure 1.**
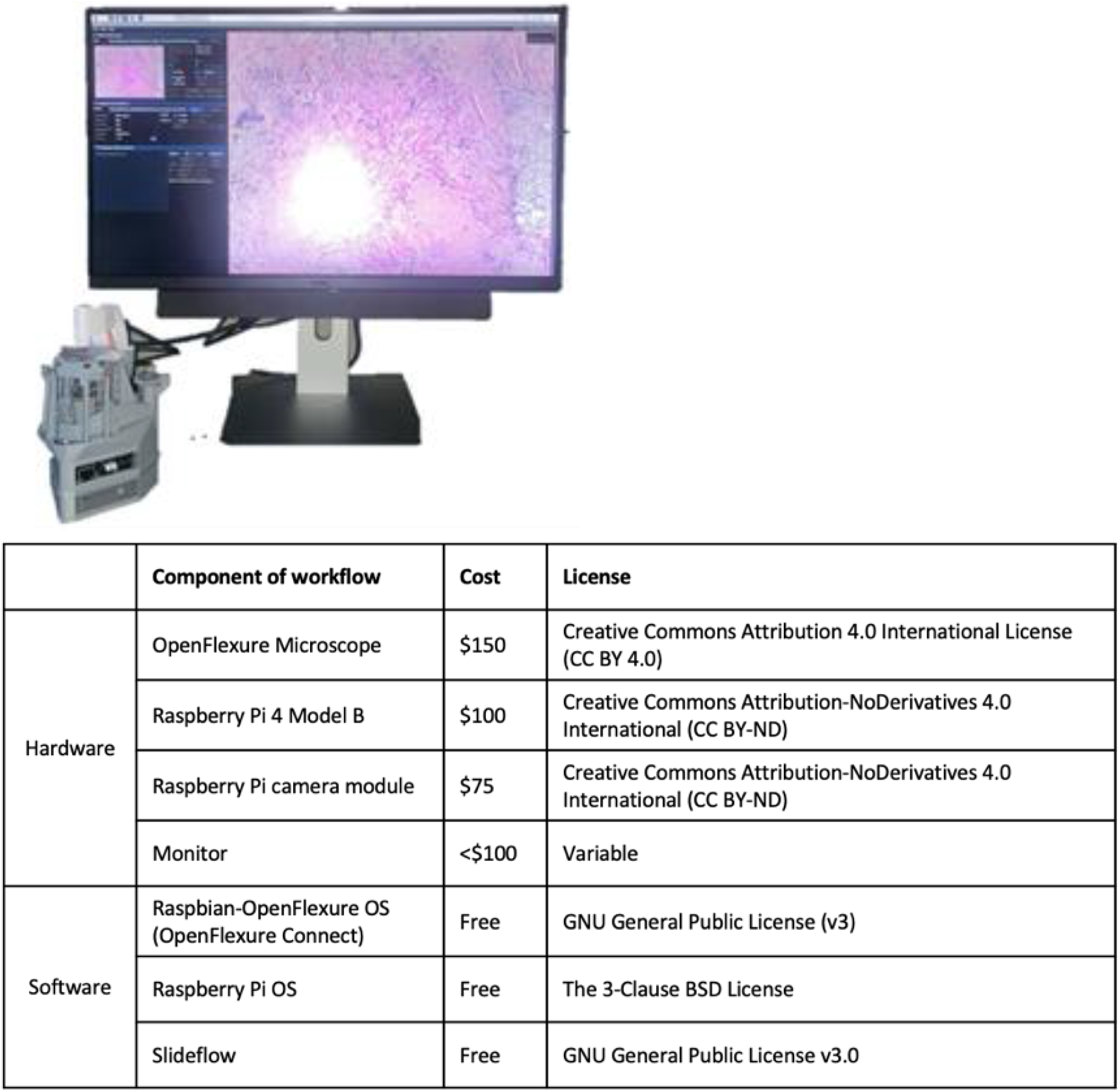
Low-cost, open-source digital pathology workstation components. Left: Assembled hardware and software components with projected digital pathology content running Slideflow software on Raspberry Pi computer. Below: Hardware and software components with respective costs and licenses.

**Figure 2.**
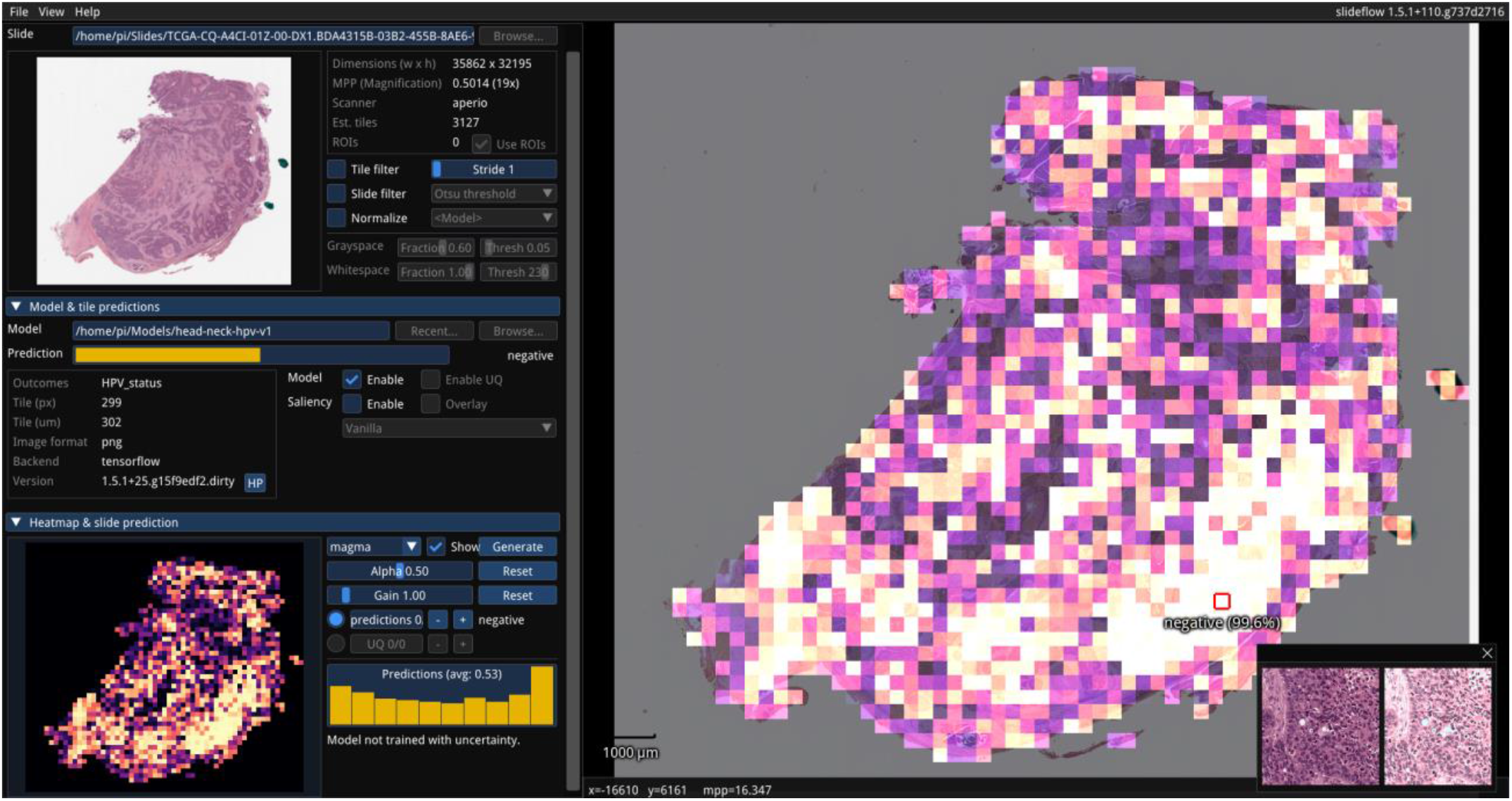
Open-source user interface for interactive visualization and generation of model predictions. Slideflow can be used to deploy a variety of trained models for digital pathology image classification, generating predictions for both partial-slide and whole-slide images. Predictions can be rendered for whole slides (rendered as a heatmap, as shown) or focal areas (rendered as individual tiles, as shown in the bottom right corner). The Slideflow user interface has been optimized for both x86 and low-power ARM-based devices. The above screenshot displays a heatmap of a WSI prediction, captured on the Raspberry Pi 4B.

The user interface utilizes a Python wrapper of Dear Imgui [25] for GUI rendering. The interface supports displaying and navigating both partial-slide and whole-slide images (JPEG, SVS, NDPI, MRXS, TIFF) with panning and zooming. Slide images are read using VIPS [26] and rendered using OpenGL 2.1+. Loading and navigating a whole-slide image utilizes <2 GB of RAM and provides a smooth experience, rendering slides at an average of 18 frames per second (FPS) while actively panning and zooming. Models trained in both PyTorch and Tensorflow can be loaded, allowing focal predictions of specific areas of a slide or whole-slide predictions and heatmaps of an entire image. All necessary preprocessing, including optional stain normalization, is encoded in model metadata and performed on-the-fly. Predictions can also be generated for WSIs, rendering a final slide-level prediction and displaying tile-level predictions as a heatmap. We implemented and optimized a low-memory mode for this interface to support whole-slide predictions on Raspberry Pi hardware. In low memory mode, partial-slide and whole-slide predictions are generated using a batch size of 4 without multiprocessing. Active CPU cooling is recommended for systems generating WSI predictions due to high thermal load associated with persistent CPU utilization.

### Statistical analysis

Deep learning model performance was assessed at patient- and tile-level using confusion matrices and area under the receiver operating curve (AUROC). For each classification model, linear regression was performed and an R^2^ value was calculated to assess the correlation between predictions made on images captured with OpenFlexure versus those captured with 3DHISTECH.

## Results

### Benchmarking DL predictions on the Raspberry Pi

We benchmarked deep learning inference speed on the OpenFlexure microscope using 24 model architectures, four different tile sizes (71×71, 128×128, 256×256, and 299×299 pixels), and a variety of batch sizes (**Supplementary Table 2**). Deep learning benchmarks were performed using the Tensorflow backend of Slideflow. Our frequently utilized, standard classification architecture (Xception at 299×299 pixels) allowed predictions at 1.04 images / second. At all tile sizes, the fastest architecture was MobileNet, allowing 4.64 img/sec at 299×299 pixels, 6.30 img/sec at 256×256 pixels, 16.72 img/sec at 128×128 pixels, and 28.50 img/sec at 71×71 pixels.

Using the whole-slide user interface, focal predictions from a model trained on 299 × 299 pixel images could be generated at approximately 1 image per second, utilizing 2.2 GB of RAM on average. WSI predictions at 10x magnification required an average of 15 minutes for a typical slide with ∼0.8 cm^2^ tumor area and utilized all 4 GB of RAM. The swap file size needed to be increased to 1 GB to generate WSI predictions. Thermal throttling was observed during WSI predictions when using standard passive CPU cooling.

### HNSCC HPV status model performance with open-source pipeline

When comparing predictions made on images from the UCMC external validation cohort captured by the 3DHISTECH versus OpenFlexure microscopes, the HNSCC HPV status model predictions had an R^2^ of 0.97 (**Figure 3**). **Figure 4** shows confusion matrices of predictions made when images were captured using the 3DHISTECH or the OpenFlexure device. With a sample size of ten patients, the AUROC for the model tested with images captured by 3DHISTECH and OpenFlexure microscopes were 0.92 and 1.00 at the patient-level and 0.71 and 0.75 at the tile-level, respectively **(Figure 5**).

**Figure 3:**
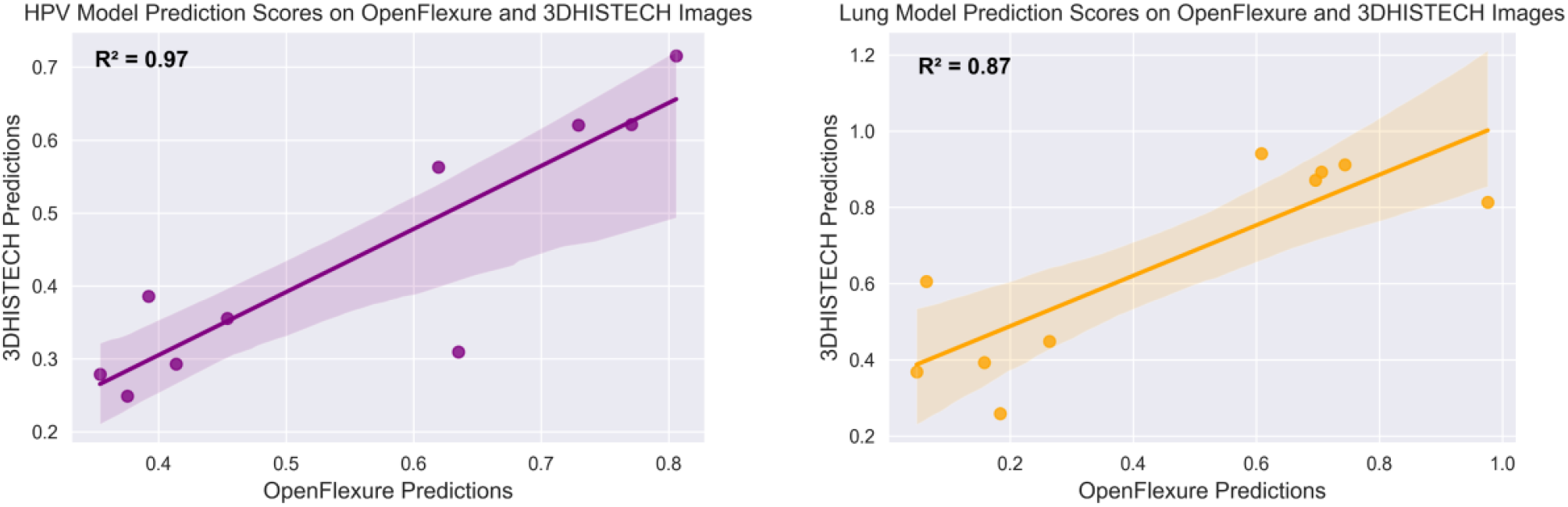
Correlation between DL numerical predictions made on images captured by 3DHISTECH vs OpenFlexure device.

**Figure 4:**
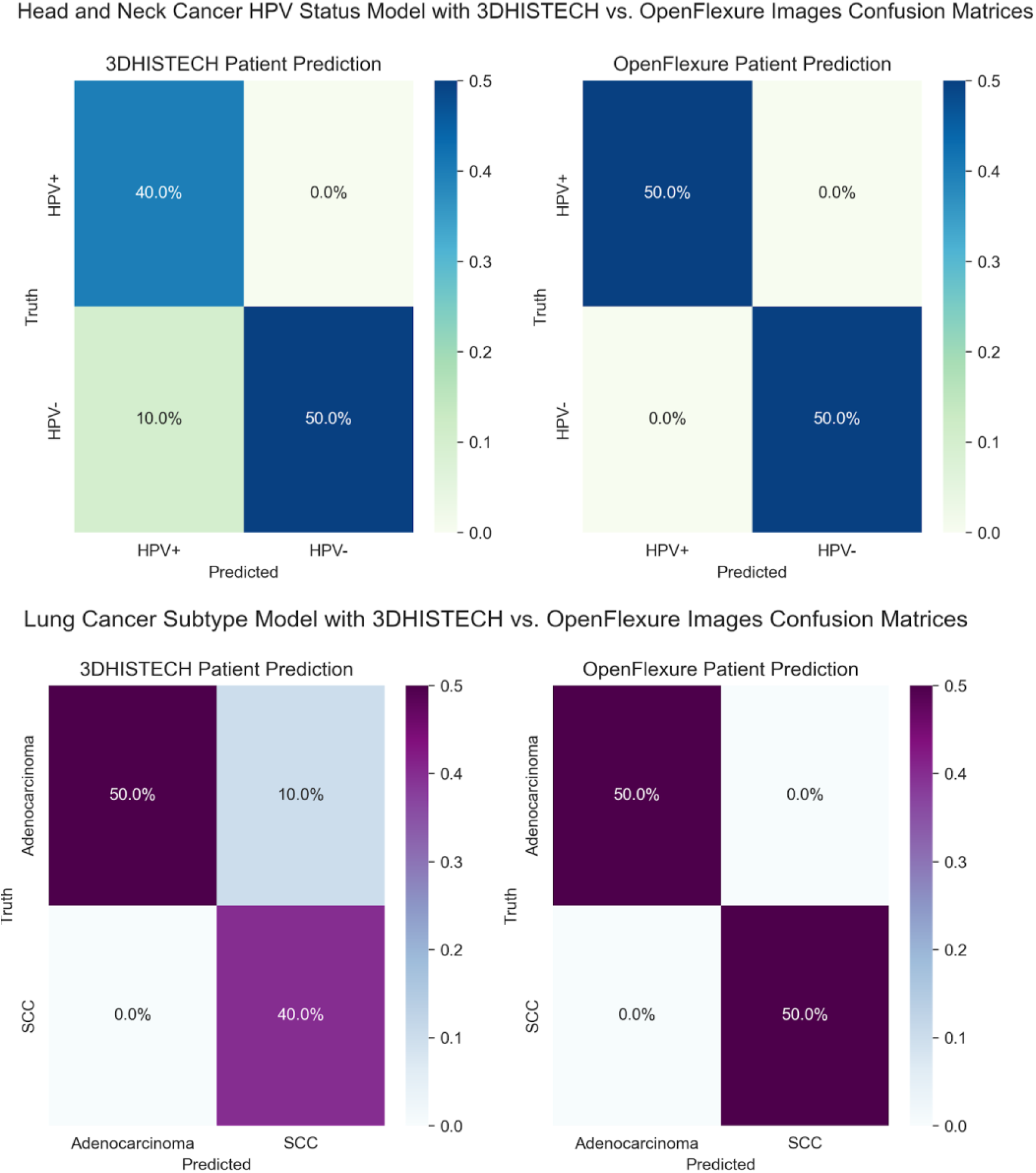
Confusion matrices showing patient-level accuracy when predictions were made using images captured with the 3DHISTECH or OpenFlexure device.

**Figure 5:**
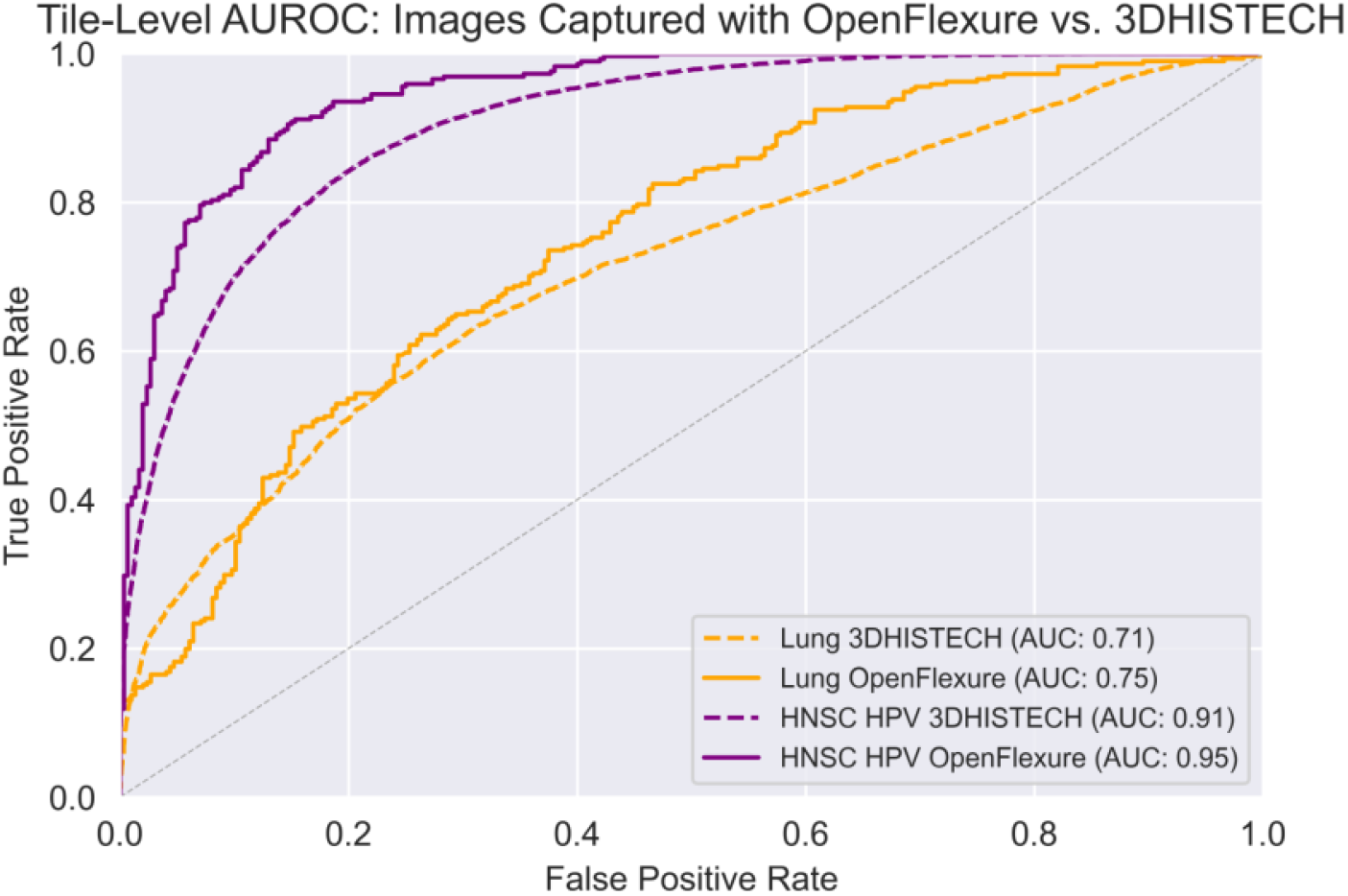
AUROC compares tile-level accuracy between high-cost 3DHISTECH and low-cost OpenFlexure image acquisition hardware.

### Lung cancer subtype model performance with open-source pipeline

Predictions made on images from the UCMC external validation cohort captured by the 3DHISTECH and the OpenFlexure microscopes had an R^2^ of 0.87 (**Figure 3**). Confusion matrices of model predictions on images captured with 3DHISTECH and OpenFlexure are shown in **Figure 4**. The patient-level and tile-level AUROCs for the model tested on images captured with 3DHISTECH were 1.0 and 0.91, respectively. The model tested on images captured by OpenFlexure performed with a patient-level and tile-level AUROC of 1.0 and 0.95 (**Figure 5**).

## Discussion

Open-source technology is key to ensuring access of artificial intelligence-based diagnostic tools to a wider range of providers and communities. It also allows users globally to contribute to the development of the clinical tools that serve the needs of their patient populations. Our results serve as proof-of-concept that it is possible to preserve model accuracy while dramatically reducing the cost of image acquisition and computational hardware. We show the feasibility of using low-cost, open-source hardware for the image acquisition and computational steps required to apply machine learning methods to digital pathology and cancer diagnostics.

Model performance for both HPV and lung classification was maintained with an open source, 3D-printed device for image acquisition despite it costing orders of magnitude less than the clinical-grade microscopes currently used. Further studies should study the influence of hardware on ML predictive accuracy in digital pathology to find more opportunities to reduce cost while maintaining quality. This work confirms the need to study the influence of image acquisition hardware on model performance before ML methods can be applied clinically.

Despite the presence of color distortion and blur prior to image normalization and lower resolution, images captured using the low-cost OpenFlexure microscope and the Raspberry Pi camera were classified by our DL model with equal accuracy as images captured on a clinical grade microscope. Model accuracy was maintained whether the task involved predicting HPV status or lung cancer subtype, supporting the idea that significant reduction in image quality associated with reduced hardware costs does not hinder model performance and can be applied to histopathologic cancer diagnostics more broadly.

Model accuracy can be further optimized by exploring strategies to computationally augment lower-quality images or to modify the training dataset such that the model can be trained on images whose quality reflects those captured by low-cost devices could improve model performance [11]. Given that stain normalization appears to have contributed to the success of the OpenFlexure device, it will be useful to explore how model performance changes without stain normalization as well as with variations in stain normalization methods. Slideflow computationally alters the training dataset by introducing random blur and JPEG compression augmentation. It will also be important to determine the influence of these techniques on model performance.

There are several limitations to this study, including the small quantity of validation data, the limited number of models tested, and the intentional use of highly informative regions of tissue for image capture to test the OpenFlexure device. Additionally, we have not accounted for any potential site/batch effects or issues including out of distribution data and domain shift [27, 28], we have not explored whether higher-level features often used as clinical biomarkers, such as breast cancer receptor status and microsatellite instability, can be identified and used to make accurate predictions with lower-resolution images such as those captured by the OpenFlexure device [12]. The TCGA data used for model training comes primarily from patients in the United States and Europe, which may limit model performance on populations with a wider range of demographics. It is crucial that DL models are trained on datasets that are representative of the intended patient populations [14]. Further studies concerned with the implementation of DL technologies in low-resource settings should explore how dataset demographics affect model performance prior to the deployment of these technologies in clinical settings.

Design improvements in the OpenFlexure microscope to improve image quality, reduce costs, and ease construction and transport are ongoing. The next iteration of the device will include an automated mechanism for whole slide image capture to ease the process of preparing scanned images for classification by the DL model. In addition, parts of the device are currently 3D printed, which might limit availability and ease of assembly.

Although this work is limited in scope, the results are promising. Further validation of existing DL models on future iterations of the OpenFlexure device and testing prediction of higher-level phenotypes, grading, and treatment response is essential to developing novel, low-cost equipment and methods for image acquisition and analysis in digital pathology that can be used to increase access to precision cancer care in lower-resource clinical settings.

