## Supplemental Tables for "Validating a low-cost, open-source, locally manufactured workstation and computational pipeline for automated histopathology evaluation using deep learning"

**Supplementary Table 1. Model parameters and hyperparameters.**

| **Hyperparameter** | **HPV** | **Lung** |
| --- | --- | --- |
| Augmentation | Flip, Rotate, JPEG Compression, Blur | Flip, Rotate, JPEG Compression, Blur |
| Batch Size | 64 | 128 |
| Dropout | 0.1 | 0.1 |
| Early Stopping | Yes | Yes |
| Early Stopping Method | Accuracy | Accuracy |
| Early Stop Patience | 0 | 0 |
| Epochs | 1 | 1 |
| Hidden Layer Width | 1024 | 1024 |
| Hidden Layers | 2 | 2 |
| L1 Regularization | 0 | 0 |
| L1 Regularization (Dense Layers) | 0 | 0 |
| L2 Regularization | 0 | 0 |
| L2 Regularization (Dense Layers) | 0 | 0 |
| Learning Rate | 0.0001 | 0.0001 |
| Learning Rate Decay | 0.97 | 0.98 |
| Learning Rate Decay Steps | 512 | 512 |
| Loss | Sparse Categorical Cross Entropy | Sparse Categorical Cross Entropy |
| Normalizer | Reinhard Fast | Reinhard Fast |
| Optimizer | Adam | Adam |
| Pooling | Average | Average |

|  | **Tile size = 71 px** | | **Tile size = 128 px** | | **Tile size = 256 px** | | **Tile size = 299 px** | |
| --- | --- | --- | --- | --- | --- | --- | --- | --- |
| **Architecture** | **Batch size = 1** | **Batch size = 8** | **Batch size = 1** | **Batch size = 8** | **Batch size = 1** | **Batch size = 8** | **Batch size = 1** | **Batch size = 8** |
| Xception | 2.15 | 8.58 | 1.89 | 4.64 | 0.91 | 1.45 | 0.74 | 1.04 |
| VGG16 | 3.13 | 6.60 | 1.49 | 2.11 | 0.45 | 0.54 | 0.35 | 0.40 |
| VGG19 | 2.67 | 5.24 | 1.19 | 1.67 | 0.37 | 0.41 | 0.28 | 0.32 |
| ResNet50 | 2.33 | 8.32 | 1.71 | 4.58 | 0.87 | 1.51 | 0.68 | 1.05 |
| ResNet101 | 1.22 | 4.50 | 0.92 | 2.44 | 0.48 | 0.80 | 0.40 | 0.57 |
| ResNet152 | 0.82 | 3.07 | 0.63 | 1.62 | 0.33 | 0.56 | 0.27 | 0.39 |
| ResNet50_v2 | 2.40 | 8.79 | 1.83 | 4.94 | 1.00 | 1.65 | 0.81 | 1.19 |
| ResNet101_v2 | 1.18 | 4.54 | 0.92 | 2.51 | 0.51 | 0.90 | 0.42 | 0.64 |
| ResNet152_v2 | 0.78 | 3.09 | 0.62 | 1.70 | 0.34 | 0.60 | 0.28 | 0.43 |
| Inception-V3 | NA | NA | 1.67 | 6.65 | 1.03 | 2.18 | 0.84 | 1.49 |
| NASNetLarge | 0.47 | 1.92 | 0.38 | 1.25 | 0.22 | 0.44 | 0.19 | 0.33 |
| Inception-ResNet-V2 | NA | NA | 0.70 | 2.70 | 0.42 | 0.90 | 0.34 | 0.65 |
| MobileNet | 6.24 | 28.50 | 4.90 | 16.72 | 3.14 | 6.30 | 2.66 | 4.64 |
| MobileNet-V2 | 4.17 | 20.52 | 3.47 | 14.07 | 2.35 | 5.53 | 2.12 | 4.18 |
| DenseNet-121 | 1.68 | 7.20 | 1.19 | 4.13 | 0.76 | 1.61 | 0.66 | 1.26 |
| DenseNet-169 | 1.24 | 5.54 | 0.90 | 3.23 | 0.57 | 1.32 | 0.52 | 1.03 |
| DenseNet-201 | 1.03 | 4.54 | 0.73 | 2.58 | 0.46 | 1.03 | 0.42 | 0.83 |
| EfficientNet-V2B0 | 2.22 | 10.72 | 1.95 | 7.91 | 1.23 | 3.89 | 1.09 | 2.93 |
| EfficientNet-V2B1 | 1.81 | 8.83 | 1.59 | 6.49 | 1.08 | 3.00 | 0.95 | 2.32 |
| EfficientNet-V2B2 | 1.73 | 8.14 | 1.47 | 5.99 | 1.00 | 2.74 | 0.89 | 2.11 |
| EfficientNet-V2B3 | 1.42 | 6.69 | 1.21 | 4.82 | 0.82 | 2.18 | 0.72 | 1.65 |
| EfficientNet-V2S | 1.10 | 4.89 | 0.90 | 3.42 | 0.60 | 1.45 | 0.51 | 1.06 |
| EfficientNet-V2M | 0.68 | 2.92 | 0.57 | 2.06 | 0.37 | 0.83 | 0.28 | 0.62 |
| EfficientNet-V2L | 0.43 | 1.76 | 0.36 | 1.18 | 0.22 | 0.42 | 0.18 | 0.31 |

**Supplementary Table 2. Deep learning inference benchmarks on the Raspberry Pi 4B.** All numbers shown indicate the speed of inference in images per second. “NA” indicates that the model could not be run due to incompatible input image size.
